# Integrin α6 upregulation in nucleus pulposus cell under high oxygen tension attenuates intervertebral disc degeneration

**DOI:** 10.1101/2021.03.10.434904

**Authors:** Zeng Xu, Jiancheng Zheng, Ying Zhang, Huiqiao Wu, Bin Sun, Ke Zhang, Jianxi Wang, Fazhi Zang, Xingkai Zhang, Lei Guo, Xiaodong Wu

## Abstract

The destruction of low oxygen microenvironment played critical roles in the pathogenesis of intervertebral disk degeneration (IVDD). In this study, high oxygen tension (HOT) treatment upregulated integrin α6(ITG α6) expression, which could be alleviated by blocking PI3K/AKT signaling pathway. And the levels of ITG α6 expression were increased in the NP tissue from IVDD patients and IVDD rat model with mild degeneration, which were reduced as degeneration degree increases. Further studies found that ITG α6 could protect NP cells against HOT-induced apoptosis and oxidative stress, and protect NP cells from HOT-inhibited ECM proteins synthesis. ITG α6 upregulation by HOT contributed to maintain a NP tissue homeostasis through the interaction with hypoxia inducible factor-1α (HIF-1α). Furthermore, silencing of ITG α6 in vivo could obviously accelerate puncture-induced IVDD. Taken together, ITG α6 upregulation by HOT in NP cells might be a protective factor in IVDD as well as restore NP cell function.

## Introduction

Intervertebral disk degeneration (IVDD) is the most common musculoskeletal disorder and a major cause of disability^1–3^. That development of IVDD is directly associated with several factors, such as aging, genetic susceptibility, body weight, heavy load work and smoking^4–5^. Intervertebral disc (IVD) mainly consist of three tissue compartments: the outer annulus fibrosus (AF), the inner nucleus pulposus (NP), and the cephalic and cauda cartilage endplates^6^. NP and AF tissues work together to provide the major anatomic functions of the IVD: spinal flexibility and load distribution^7^. NP cells play the core role in the maintenance of IVD function^6^. The number and function of NP cells are decreased and the secretion of inflammatory factors is increased with increasing age in human, which is characterized by increased matrix catabolism, fibrosis accompanied by changes in NP cell phenotype^8–9^.

IVD is the largest ischemic and hypoxic tissue in mammals, due to a complete lack of tissue vascularity^10^. Thus, NP cells are characterized by low oxygen tension. Hypoxia plays important roles in maintaining the physiological functions of NP cells, such as energy metabolism, matrix synthesis, and cell viability^11–12^. NP cells could ensure their function and survival in low oxygen environment through the regulation of cells apoptosis, the generation of ROS^13^ and some critical genes expressions, such Sox9 and Collagen II alpha 1(Col2α1). Growing evidences suggested that neovascularization in discs increased oxygen tension in the progression of IVDD, which lead to perturbation of low oxygen gradient and the change of cell metabolism^14–15^. However, the exact mechanisms by which the change of low oxygen microenvironment initiates IVDD is not fully elucidated.

Integrin (ITG), which are well-known receptors, are important in the communication between cells and its surrounding tissues or extracellular matrix (ECM) to regulate cell apoptosis, migration, proliferation and survival^16^. Integrins are comprised of two distinct chains: alpha (α) subunits and beta (β) subunits. α and β subunits have been discovered in vertebrates, which totally assemble into 24 distinct integrin complexes^17^.Increasing evidences suggested that integrins were involved in the pathogenesis of IVDD^18–20^. As compared to adjacent annulus fibrosus region, several integrin subunits were more highly expressed in nucleus pulposus cells, such as ITG β4,α3, α5,and α6^21^.Recently studies showed that ITG α6 was involved in the regulation of the production of ROS and cell apoptosis by hypoxia in some organs and tissues^22–23^. However, it is unclear whether ITG α6 contributes to the regulation of NP cells by change of the oxygen content, especially when exposed to high oxygen tension (HOT). In this study, we found that the levels of ITG α6 expression were increased in the NP tissue from IVDD patients and IVDD rat model with mild degeneration, which were reduced as the degree of degeneration increases in severity. ITG α6 expression was also significantly up-regulated by HOT through activation of PI3K/AKT signaling pathway in NP cells. Inhibition of hypoxia inducible factor-1α (HIF-1α) by HOT could increase the expression of ITG α6, whereas increasing of ITG α6 level downregulated HIF-1α expression. Further studies showed that ITG α6 was involved in the regulation of NP cells functions, including NP cells apoptosis, the Reactive Oxygen Species (ROS) production and ECM secretion. In addition, silencing of ITG α6 accelerated puncture-induced IVDD *in vivo*. Taken together, these data indicate that ITG α6 may be a protection factor for the pathogenesis of IVDD.

## Methods and materials

### Reagents and antibodies

Fetal bovine serum (FBS) was purchased from Biological Industries (Israel). Phosphate buffered solution (PBS) and Dulbecco’s modified Eagle’s medium/F12 (DMEM/F12) were purchased from Hyclone (South Logan, UT, USA).Improved Minimal Essential Medium(Opti-MEM), 0.25% trypsin/ethylene diamine tetraacetic acid(EDTA) and type II collagenase were purchased from Gibco(Gaithersburg, MD, USA). Trizol and Lipofectamine 3000 was purchased from Invitrogen (Carlsbad, CA, USA). The ITG α6-specific siRNA duplexes were designed previously and synthesized by RiboBio (Guangzhou, China). PrimerScript RT Master Mix and PrimerScript RT Master Mix were purchased from Takara Bio Inc (Shiga, Japan).The JC-1 assay kit, TUNEL Apoptosis Assay Kit, Reactive Oxygen Species(ROS) assay kit, RIPA buffer, Annexin V-FITC/propidium iodide (PI) staining kit, PI3K inhibitor(LY294002), the antibody against HIF-1α, Bax and caspase3 were all purchased from Beyotime Biotechnology (Shanghai, China). The antibodies against Col2α1, Aggrecan and β-actin were purchased from Abcam (Cambridge, MA, USA). The antibody against ITG α6 was purchased from Sabbiotech (College Park, Maryland, USA). The antibody against AIF was purchased from Cell Signaling Technology (Danvers, MA, USA). The antibody against AKT, p-AKT, BCL-2 were purchased from ABclonal (Woburn, MA, USA).Chitosan hydrochloride was purchased from Ultrapure Aoxing Bio(Zhejiang, China). Pentasodium tripolyphosphate(TPP) was purchased from Sigma (St. Louis, MO, USA).The HIF-1α knockout (HIF-1α ^-/-^) mouse model was donated by Professor Zhang.

### Human IVD specimen collection

The degeneration degree of IVDD was assessed according to the Pfirrmann classification system(I-V) ^24^, which entails the use of T2-weighted magnetic resonance imaging (MRI). To minimize the errors as much as possible, every specimen was assessed by three independent observers. Degenerated specimens were obtained from patients who underwent spinal fusion surgery with low back pain (LBP), normal discs specimens were obtained from young patients with burst lumbar fracture without IVDD. Finally, twenty-five specimens were collected including 4 healthy discs (19-27 years old, two males and two females) and 21 various degeneration degree discs (22-56 years old, nine males and twelve females). The specimens were divided into four groups: normal (Pfirrmann I, *n=4*), mild degeneration (Pfirrmann II, *n=5*), moderate degeneration (Pfirrmann III, *n=7*), severe degeneration (Pfirrmann IV-V, *n=9*).

### Establishment of IVDD model

The animals used in this study were 3-month-old male Sprague-Dawley(SD) rats, weighing 200–250g.The annulus needle puncture model of IVDD in SD rat caudal spine was similar to the previous study^25^. Totally eighteen rats were randomized as follows: control, 2-and 4-weeks groups with six rats in each group. After anesthesia with intramuscular injection of 1% pentobarbital sodium at 0.1 mg/kg, a portable X-ray machine was used to check the coccygeal 4-5disc level (Co4-5), which is the surgical level. A 28-gauge needle was inserted into the center of the disc through the AF, then rotated 360° and held for 30s. The rats were euthanized and the Co4-5 caudal disc were collected. Nine specimens were stored at −80°C for protein extraction for western blotting. The remaining specimens were fixed, decalcified, paraffin-embedded and sectioned, then were stained with hematoxylin and eosin (HE), Alcian Blue and immunofluorescence for analysis.

### Histology staining

Both human and rat specimens were fixed with 4% formaldehyde, rat IVD tissues also need to be decalcified with 10% ethylenediaminetetraacetic acid solution for 4 weeks. Then all specimens were paraffin-embedded for sectioning longitudinally in the sagittal plane at 5-mm intervals. Sections were stained with HE and Alcian Blue to analyze the morphological changes in the discs.

### Tissues and cells immunofluorescence staining

Immunofluorescence to assess ITG α6, HIF-1α, AIF, Col2α1 expression was performed on sections of IVD tissues and fixed cell samples. The fixed cells or sections were blocked with 3% BSA for 1 h, then washed with PBS. Subsequently, samples were incubated overnight with corresponding antibody mentioned above at 4°C. After three washes with PBS, samples were incubated with a goat anti-rabbit or mouse IgG antibody (1:5000, 2h, room temperature). The samples were then incubated with 0.1 g/mL DAPI for 3 min. After the final round of washes, fluorescence images were captured under a fluorescence microscope microscope (LSM 710, Carl Zeiss, Oberkochen, Germany).

### NP cells isolation and culture

NP cells were isolated and cultured as previously described^26^. Briefly, the SD rats were killed with over-dose anesthesia. The caudal vertebra spine was totally taken out. Under aseptic condition, the skin and the musculature attached to the vertebral column were removed to exposed the IVD. Then, he annulus fibrosus was incised with bistoury to expose and separate the nucleus pulposus tissues. After cutting the NP tissues into pieces, NP tissues was subsequently digested with 0.25 % trypsin with EDTA for 15 mins and 0.1% type II collagenase for 2 h at 37 °C. NP cells were obtained after filtration of the above residual tissues. Then the suspension was centrifuged at 1500rpm for 5mins.The cell pellet was resuspended in DMEM/F12 contained 10% FBS and seeded into 10cm plate in a humidified atmosphere of 1% O_2_, 5% CO_2_, at 37 °C. NP cells were adherent to the bottom of culture dish after 3 days. Cell growth was observed every day under a microscope. Culture medium was replaced by fresh one for the first time on the 6th day and every 3 days. When confluent after 8-10 days, NP cells were lifted using 0.25 % trypsin with EDTA and subcultured to the second generation. And the third generation were used for the following experiments.

### Cell transfection

Lipofectamine 3000 reagent was used to transiently deliver ITG α6 siRNA or negative control (NC) siRNA into NP cells according to the protocol. Briefly, ITG α6 or NC siRNA was mixed with Lipofectamine 3000 reagent and Opti-MEM at room temperature for 15 mins. The mixture was added into cells at 90% confluence. After culturing at 37°C and 1% O_2_ for 24h, total protein was extracted and analyzed by western blot or qRT-PCR to demonstrate the efficiency of transfection.

### qRT-PCR

Total RNA was extracted from cell samples using Trizol according to the manufacturer’s instructions and total RNA was measured spectrophotometrically at 260 nm. mRNA expression was detected by qRT-PCR as previously reported^27^. All data were subsequently normalized to the β-actin mRNA level and expressed as relative mRNA expression.

### Western blot

NP tissues separated from the IVD was firstly ground to powder via the grinder. Then NP cells or NP tissues were lysed in RIPA buffer containing protease and phosphatase inhibitor for 30 mins at 4°C. The supernatant was carefully collected in order to acquire total protein. Protein concentration was measured by BCA protein assay kit. The samples were electrophoresed on SDS–PAGE and transferred to PVDF membranes. The membranes were blocked with 5% nonfat milk for 30min to block non-specific antigens followed by corresponding primary antibodies and incubated overnight at 4 °C. After washing in TBST 3 times for 10 min, the membrane was incubated with a secondary anti-IgG-HRP antibody at room temperature for 2 h. Immunolabeling was detected using enhanced chemiluminescence reagent.

### Flow cytometry analysis for apoptosis percentage and ROS production level

The apoptosis percentage of NP cells was analyzed using an Annexin V-FITC/propidium iodide (PI) staining kit, and the ROS production level was analyzed using a Reactive Oxygen Species assay kit, both experiments were performed following the manufacturer’s protocol. Briefly, after being digested with 0.25% trypsin and centrifuged (1500rpm for 5 min), NP cells was obtained and then incubated with Annexin V-FITC for 10 min, and then PI for 10 min for apoptosis percentage analysis or incubated with DCFH-DA for 20mins for ROS production level detection. Stained cells were quantified by flow cytometry.

### TUNEL

The cells treated as indicated were fixed with 4% paraform phosphate buffer saline, rinsed with PBS, then permeabilized by 0.1% Triton X-100 for FITC end-labeling the fragmented DNA of the apoptotic NP cells using TUNEL cell apoptosis detection kit. The FITC-labeled TUNEL-positive cells were imaged under a fluorescent microscopy by using 488-nm excitation and 530-nm emission.

### Mitochondrial membrane potential assay with JC-1

The mitochondrial membrane potential was detected with a Mitochondrial Membrane Potential Assay Kit. In brief, the NP cells were seeded in 6-well plates. Following different treatments, the media were removed then were stained with JC-1 for 30mins and Subsequently, the cells were washed with serum-free medium for three times, and the fluorescence intensity (excitation 585nm, emission 590nm) was detected by a fluorescent microplate reader.

### Detection of ROS production level

ROS was estimated by a Reactive Oxygen Species assay kit according to the manufacturer’s instructions. The NP cells were seeded in 6-well plates. Following different treatments, the media were removed, and the cells were loaded with DCFH-DA (10 μM) at 37°C for 20 min. Subsequently, the cells were washed with serum-free medium for three times, and the fluorescence intensity (excitation 488nm, emission 525nm) was detected by a fluorescent microplate reader.

### Chitosan nanoparticles for siRNA delivery preparation

Nanoparticles were produced based on modified ionic gelation of TPP with chitosan as previously described^28^. Briefly, chitosan hydrochloride was dissolved with distilled water to get chitosan solution (10ml, 2mg/ml), then the solution was centrifugated at 13,000×g for 10 min. The supernatants were discarded and chitosan nanoparticles were resuspended in filtered distilled water. Do the same to get TPP aqueous solution(10ml,0.84mg/ml).

Nanoparticles were spontaneously obtained upon the addition of 0.4 ml of a TPP aqueous solution (0.84 mg/ml) to 1ml of chitosan solution (2 mg/ml, at chitosan to TPP weight ratio of 6:1) under constant magnetic stirring at room temperature. For the association of siRNA with the chitosan–TPP nanoparticles (chitosan–TPP–siRNA), 20ul of siRNA (1μg/μl) in RNase-free water was added to TPP solution (0.4ml, 0.84 mg/ ml) before adding this to the chitosan solution (1ml, 2 mg/ml). The particles were then incubated at room temperature for 30 min before further use.

### Injection of chitosan-TPP-siRNA nanoparticles in rats IVDD model

All surgical procedures were similar to the protocol used in the establishment of IVDD rat tail model as described before. At last, 4ul of chitosan-TPP-siRNA nanoparticles or PBS was injected into the surgery level. Two weeks after surgery, specimens were obtained.

### Statistical analysis

Data were collected from three or more independent experiments and expressed as mean ± SD. A two-sided Student’s t-test was used to analyze the difference for experiments with two subgroups. One-way analysis of variance (ANOVA) was performed to show the difference for experiments with more than two subgroups.

## Results

### Upregulation of ITG α6 expression by HOT in NP cells through activation of PI3K/AKT signaling pathway *in vitro*

A panel of qRT-PCR assays were used to measure twelve known subunits of the ITG in NP cells exposed to HOT (21%, O_2_) for 24h. All of these subunits were differentially expressed in NP cells in response to HOT, in which ITG α5, ITG α6, ITG β1, ITG β4 and ITG β5 were upregulated, but ITG α1, ITG α2, ITG α3, ITG α4, ITG αD, ITG β2 and ITG β3 were downregulated (Figure 1A). Among these differentially expressed members, ITG α6 was found to be upregulated by more than two-folds. Further studies showed that ITG α6 expression was increased by HOT in a time-dependent manner in NP cells, as evidenced by the immunofluorescence staining (Figure 1B-C). QRT-PCR and western blot results also showed that HOT significantly increased ITG α6 expression in NP cells in a time dependent manner (Figure 1D-F). Furthermore, we found that the expression of p-AKT was increased in NP cells exposed to HOT from 12h to 48h (Figure 1E-H). Blocking PI3K/AKT signaling pathway with PI3K inhibitor could obviously alleviate the upregulation of ITG α6 expression by HOT (Figure 1I-L), indicating that HOT upregulated the expression of ITG α6 in NP cells through activation of PI3K/AKT signaling pathway.

**Figure 1.**
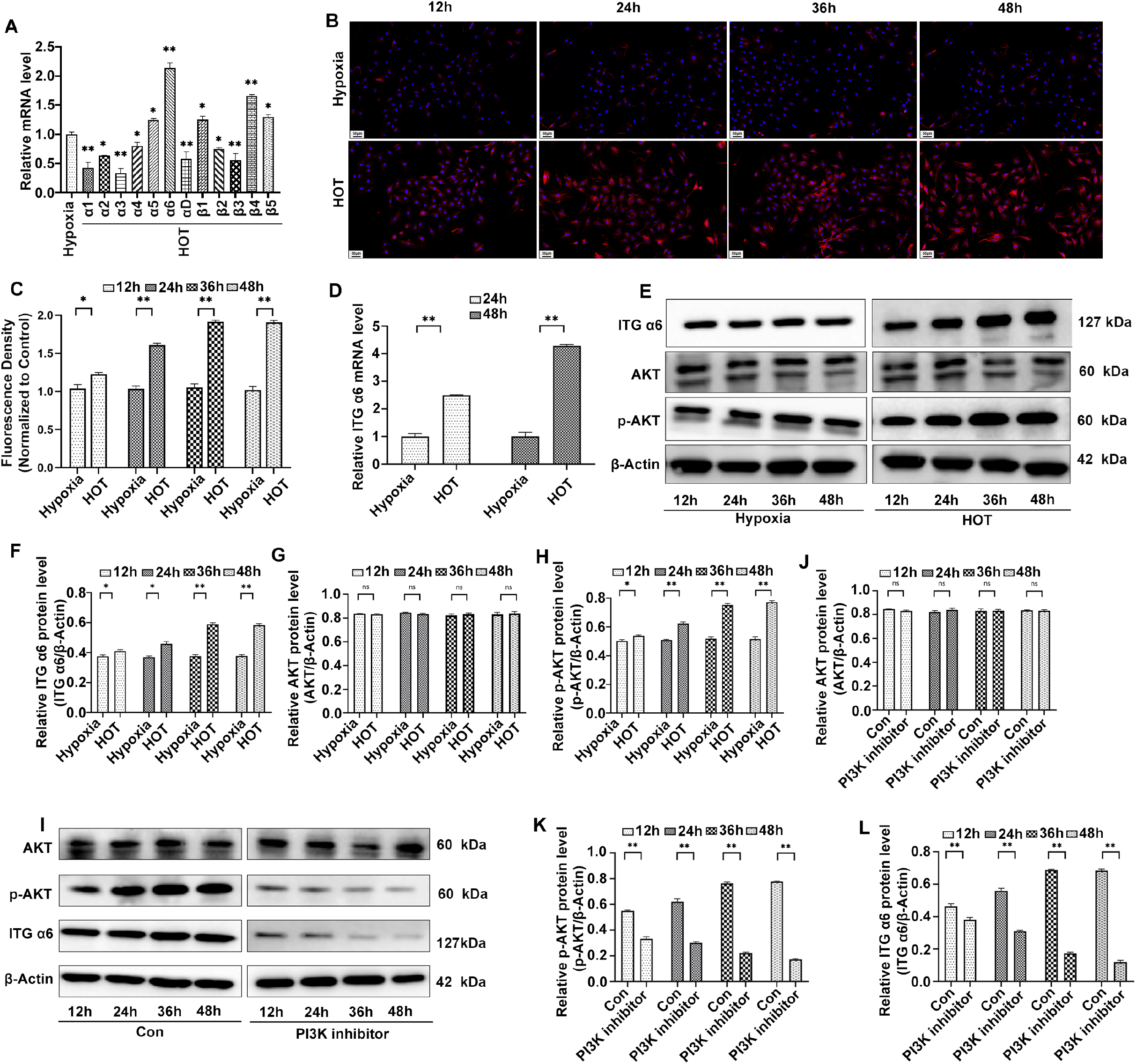
Upregulation of ITG α6 expression by HOT in NP cells through activation of PI3K/Akt signaling pathway *in vitro*. A: QRT-PCR analysis of twelve known subunits of the integrins in NP cells exposed to HOT (21%, O_2_) for 24h. *n = 4, *P < 0.05, **P < 0.01*. B: Expression of ITG α6 was observed by immunofluorescence staining in NP cells exposed to HOT (21%, O_2_) for different time course. *N = 4*. C: Quantitative analysis of positive staining. The intensity of the immunofluorescence of ITG α6 was markedly increased in NP cells exposed to HOT (21%, O2) from 12h to 48h. *n = 4, *P < 0.05, **P < 0.01*. D: QRT-PCR analysis of ITG α6 mRNA expression in NP cells exposed to HOT for 24h and 48h. *n = 4, **P < 0.01*. E: Western blot analysis of ITG α6, AKT, p-AKT and β-actin expression in NP cells exposed to HOT (21%, O_2_) for different time course. *n = 4*. F-H: The results of western blot analysis are expressed as percentages of positive mean values ± SD, *n = 3, **P < 0.01, *P < 0.05*. I: Western blot analysis of ITG α6, AKT, p-AKT and β-actin expression in NP cells treated with (LY294002,20uM) under HOT (21%, O_2_). *n = 4*. J-L: The results of western blot analysis are expressed as percentages of positive mean values ± SD, *n = 3, **P < 0.01, *P < 0.05*.

### The expression of ITG α6 is increased in the NP tissue from patients with mild degeneration of IVDD

IVDD based on MRI Data was graded by using the Pfirrmann system. A total of 25 discs NP samples were divided into four groups: normal (Pfirrmann I, *n = 4*), mild degeneration (Pfirrmann II, *n = 5*), moderate degeneration (Pfirrmann III, *n = 7*), severe degeneration (Pfirrmann IV-V, *n = 9*) (Figure 2A). HE staining and Alcian Blue staining of NP tissue sections were used to evaluate the pathologic change of the IVD tissue. The results showed that the gelatine NP gradually disappeared and fibrocartilage was formed on the surface of cartilage in the IVDD group compared to the normal group, which appeared in an IVDD degree-dependent manner (Figure 2B). Subsequently, immunofluorescence staining was applied to investigate the localization and expression of ITG α6 in NP tissue from patients with different degrees of IVDD. The results showed that the intensity of the immunofluorescence of ITG α6 was increased in the NP tissue from IVDD patients with mild degeneration. However, ITG α6 expression was decreased in human moderate and severe degenerated NP tissue (Figure 2C-D). Similar results were also examined by western blot (Figure 2E-F). These results indicated that the level of ITG α6 was increased in the early stage of disc degeneration, whereas which was reduced in the middle and late stage of disc degeneration.

**Figure 2.**
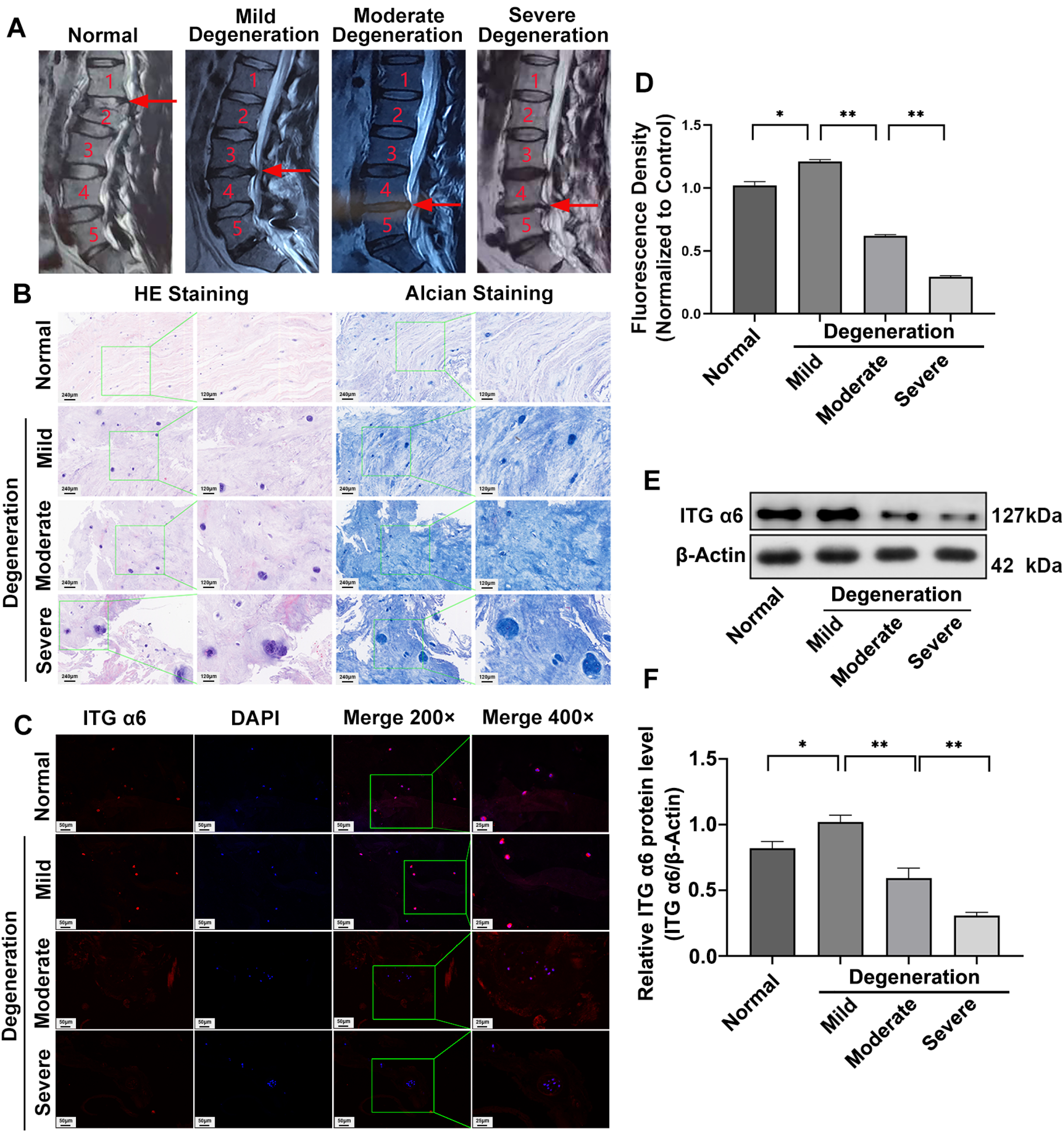
The expression of ITG α6 is increased in the NP tissue from patients with mild degeneration of IVDD. A: Representative T2-weighted MRI images of human NP tissues graded by the Pfirrmann system: normal (Pfirrmann I), mild degeneration (Pfirrmann II), moderate degeneration (Pfirrmann III), severe degeneration (Pfirrmann IV-V). B: HE staining and Alcian Blue staining of human NP tissues with different degrees of IVDD, *n = 3*. C: Immunofluorescence staining of ITG α6 in human NP tissue with different degrees of IVDD. D: Quantitative analysis of positive immunofluorescence staining. *n = 3, **P < 0.01, *P < 0.05*. E: Western blot analysis of ITG α6 in human NP tissue with different degrees of IVDD. F: The results of western blot analysis are expressed as percentages of positive mean values ± SD. *n = 3, **P < 0.01, *P < 0.05*.

### The expression of ITG α6 is increased in the NP tissue from rat model with mild degeneration of IVDD

To further examine the expression of ITG α6 in IVDD, an IVDD rat model was established by annulus needle puncture. Two and four weeks after puncture, the gelatine NP gradually disappeared and fibrocartilage was formed on the surface of cartilage, accompanied by the disappearance of the border between NP and AF, as evidenced by HE staining and Alcian Blue staining (Figure 3A-B). These results indicated that the IVDD rat model was successfully made in this study. Next, we tested the expression of ITG α6 in NP tissue from IVDD rat model at two and four weeks after puncture. The results of western blot showed that the level of ITG α6 was significantly upregulated in NP tissue from the IVDD rat model at two weeks after puncture, which was decreased in IVDD rat model at four weeks after puncture(Figure 3C-D), similar results were tested by the immunofluorescence staining (Figure 3E-F). In addition, we tested the expression of HIF-1α in NP tissue from IVDD rat model. We found that HIF-1α expression was decreased in the IVDD rat model at two weeks after puncture, which was increased at four weeks after puncture (Figure 3G-H). In addition, TUNEL staining was applied to examine the apoptosis in NP tissue from IVDD rat model. We found that the apoptotic rate of the NP cells was significantly increased in NP tissue from the IVDD rat model at two weeks after puncture, which were further increased after four weeks (Figure 3I-J).

**Figure 3.**
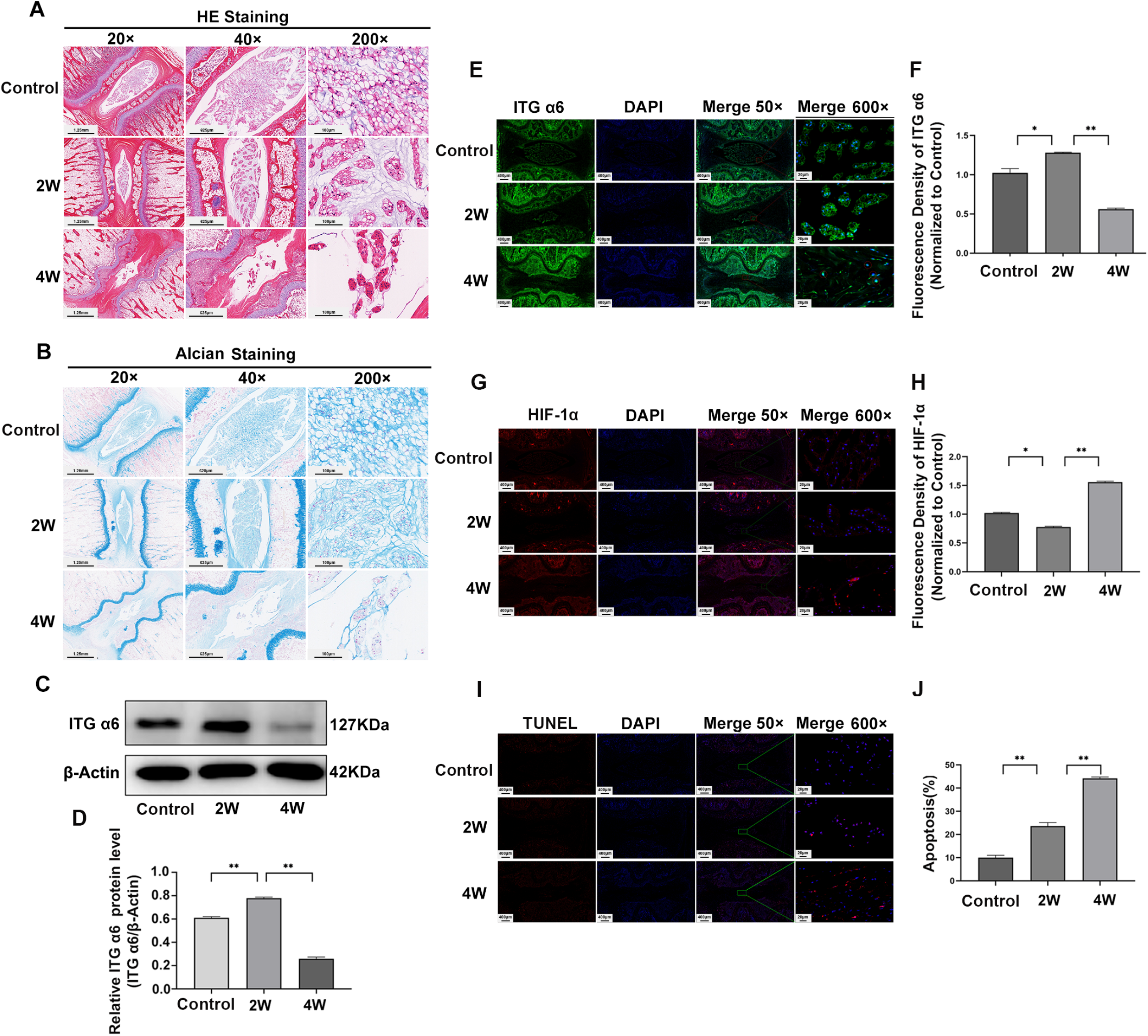
The expression of ITG α6 is increased in the NP tissue from rat model with mild degeneration of IVDD. A-B: HE staining and Alcian Blue staining of NP tissues in rat model at 2 and 4 weeks after annulus needle puncture. C: Western blot analysis of ITG α6 and β-actin expression of NP tissues in rat model. D: The results of western blot analysis are expressed as percentages of positive mean values ± SD, *n = 4, *P < 0.05, **P < 0.01*. E: Immunofluorescence staining of ITG α6 in NP tissue in rat model. F: Quantitative positive immunofluorescence staining analysis of ITG α6. *n = 4, *P < 0.05, **P < 0.01*. G: Immunofluorescence staining of HIF-1α in NP tissue in rat model. H: Quantitative positive immunofluorescence staining analysis of HIF-1α. *n = 4, *P < 0.05, **P < 0.01*. I: TUNEL staining of NP tissues to examine the apoptosis in rat model. J: Quantitative positive staining analysis of TUNEL staining. *n = 4, *P < 0.05, **P < 0.01*.

### Activation of PI3K/AKT/ITG α6 pathway protects NP cells against HOT-inhibited ECM proteins synthesis

To investigate the role of ITG α6 in the secretion of ECM, the level of ITG α6 was decreased with siRNA-ITG α6. The western blot results showed that ITG α6 expression was obviously decreased in NP cells transfected with siRNA-ITG α6 compared to NP cells transfected with siRNA-NC, indicating that the deletion of ITG α6 was effective (Figure 4A-B). The ECM consists primarily of aggregating proteoglycans and Col2α1. The western blot results showed that HOT decreased the expression of Aggrecan and Col2α1, whereas which were further decreased by silencing of ITG α6, suggesting that ITG α6 could alleviate the inhibitory effect of HOT on ECM secretion (Figure 4A-B). Furthermore, the decreasing of aggrecan and Col2α1 expression were observed in NP cells exposed to HOT from 12h to 48h, which could be aggravated by PI3K inhibitor (Figure 4C-E), suggesting that activation of PI3K/AKT/ITG α6 pathway had protective effect on NP cells against HOT-inhibited ECM proteins synthesis.

**Figure 4.**
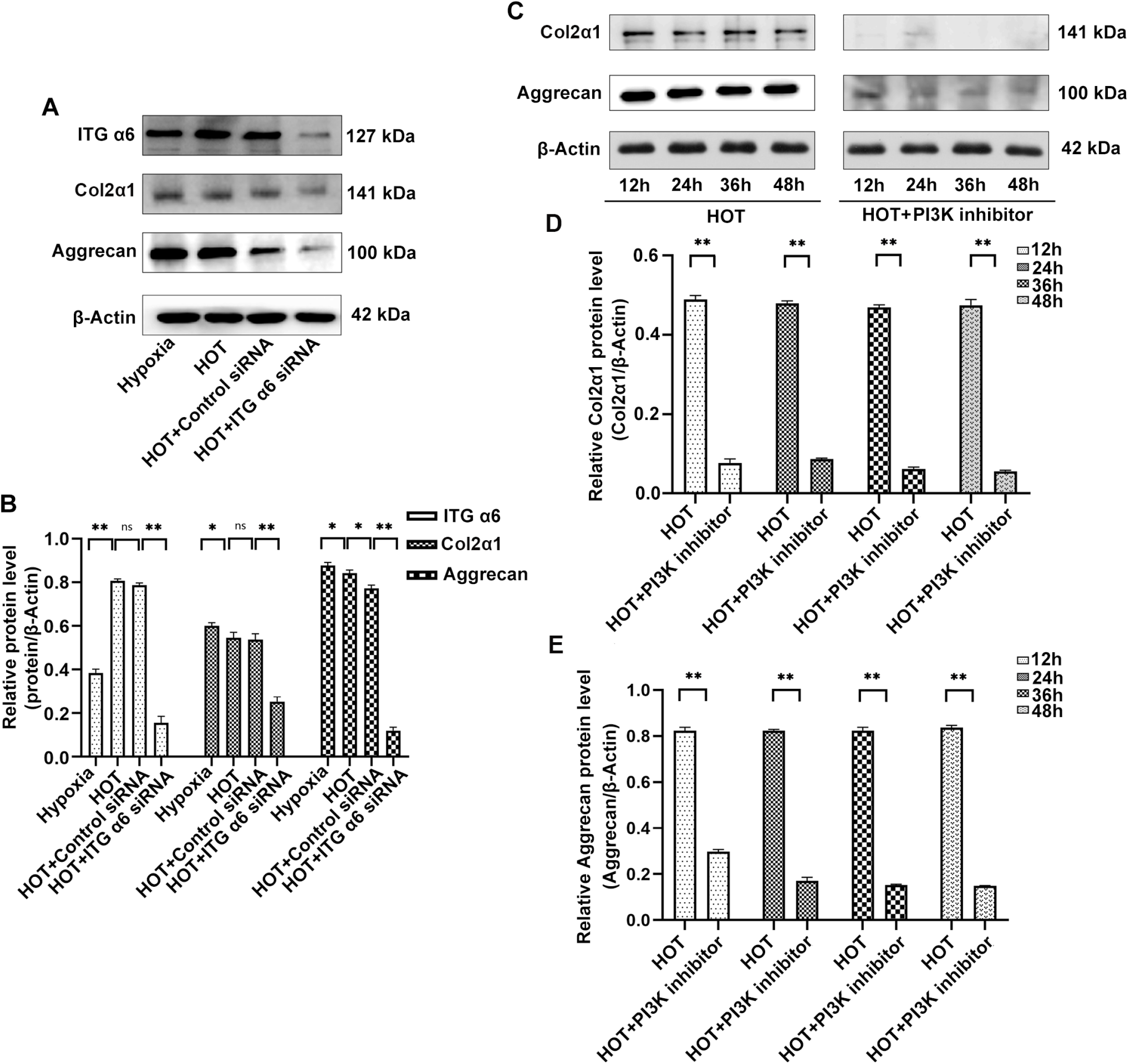
Activation of PI3K/AKT/ITG α6 pathway protects NP cells against HOT-inhibited ECM proteins synthesis. A: Western blot analysis of ITG α6, Col2α1, aggrecan and β-actin expression in NP cells treated with HOT and siRNA-ITG α6. B: The results of western blot analysis are expressed as percentages of positive mean values ± SD. n=3, \**P<0.05, **P<0.01*.C: Western blot analysis of Col2α1, aggrecan and β-actin expression in NP cells treated with HOT and PI3K inhibitor(LY294002,20uM)for 24h and 48h. D-E: The results of western blot analysis are expressed as percentages of positive mean values ± SD. *n = 3, *P < 0.05, **P < 0.01*.

### ITG α6 protects NP cells from apoptosis induced by HOT

To investigate whether ITG α6 was involved in HOT-induced NP cells apoptosis, flow cytometric analysis was firstly applied to monitor cells apoptosis. A population of apoptotic NP cells presented phosphatidylserine (PS) externalization as evidenced by a significant increase in annexin-FITC fluorescence, as well as decreased plasma membrane integrity as determined by a modest increase in propidium iodide fluorescence. We found that HOT obviously promoted serum deprivation-induced NP cells apoptosis, whereas silencing of ITG α6 further aggravated HOT-induced NP cells apoptosis (Figure 5A-B). Subsequently, the quantitative analysis of JC-1-stained cells revealed a significant increase in the green (high ^Δ^Ψm) to red (low ^Δ^Ψm) ratio in HOT-treated cells when compared with hypoxia cells. Silencing of ITG α6 could further increased green (high ^Δ^ψm) to the red (low ^Δ^ψm) to ratio in HOT-treated cells, suggesting that ITG α6 could protect NP cells from HOT-induced apoptosis (Figure 5C-D). Apoptosis inducing factor (AIF) was involved in mitochondrial respiration and caspase independent apoptosis. The results of immunofluorescence staining showed that the nuclear accumulation of AIF in NP cells was markedly increased by HOT. Silencing of ITG α6 enhanced HOT-induced the nuclear accumulation of AIF in NP cells (Figure 5E-F). The effects of silencing of ITG α6 on cells apoptosis-related proteins (BCL-2, Bax and Caspase 3) expression were observed by western blot in NP cells exposed to HOT. HOT increased the expressions of Bax and caspase 3 and decreased the expression of Bcl2, which were further aggravated in NP cells with silenced expression of ITG α6, indicating that silencing of ITG α6 further promoted HOT-induced NP cells apoptosis (Figure 5G-H). Taken together, these results indicated that ITG α6 protected NP cells from apoptosis induced by HOT.

**Figure 5.**
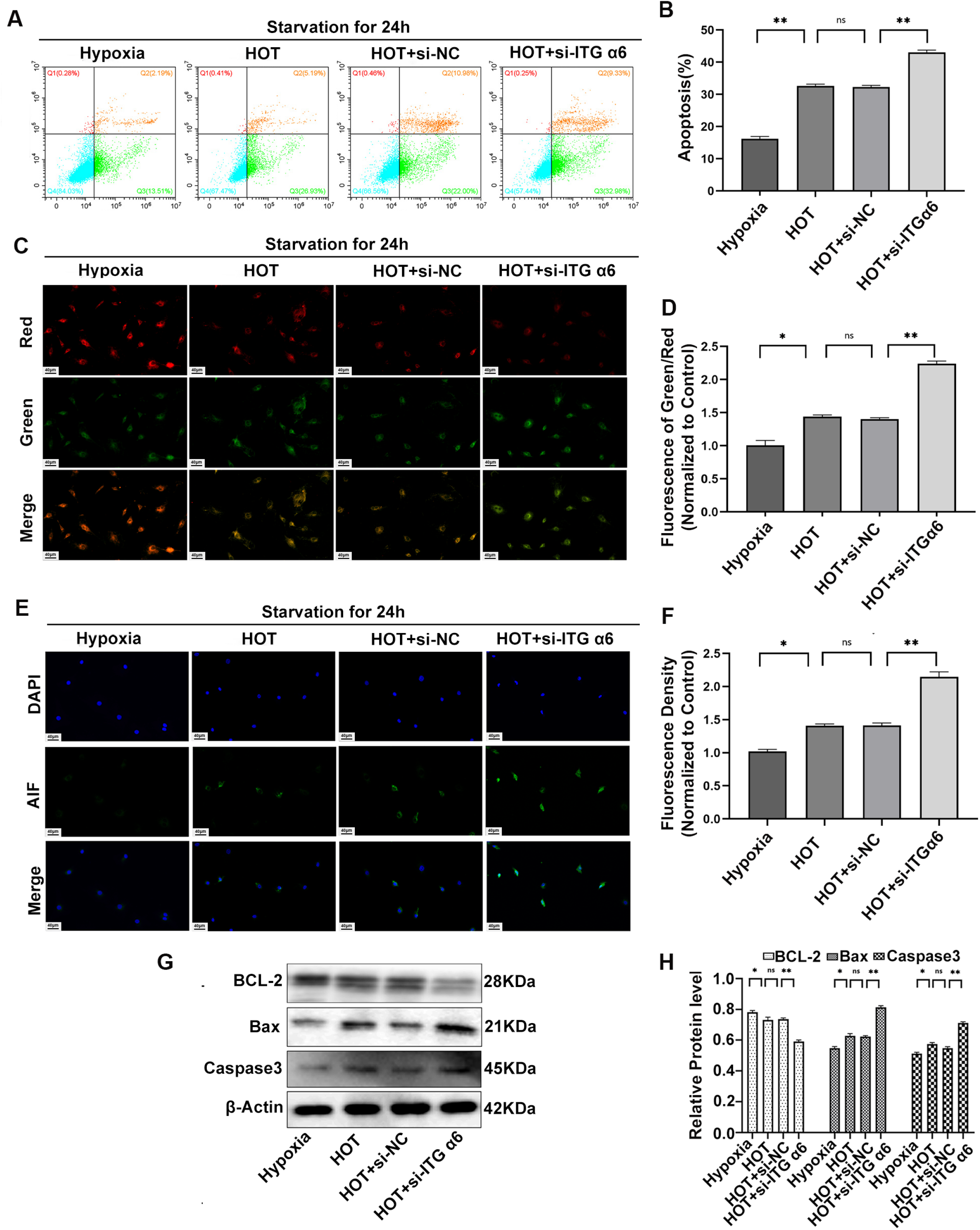
ITG α6 protects NP cells from apoptosis induced by HOT. A-B: NP cells treated with HOT and siRNA-ITG α6 under serum-free condition were stained with Annexin-V and PI and detected by flow cytometric analysis. Apoptosis percentage was calculated according to the result of flow cytometric analysis. Data were presented as the mean ± SD. *n = 3, *P < 0.05, **P < 0.01*.C-D: NP cells treated with HOT and siRNA-ITG α6 under serum-free condition were stained with JC-1. Quantitative analysis of the green (high^Δ^ψm) and red (low^Δ^ψm) staining. The green (high ^Δ^ψm) to red (low ^Δ^ψm) ratio markedly increased in NP cells treated with siRNA-ITG α6 and HOT. *n = 3, *P < 0.05, **P < 0.01*.E: Immunofluorescence staining of AIF in NP cells treated with HOT and siRNA-ITG α6 under serum-free condition. F: Quantitative analysis of positive immunofluorescence staining. *n = 3, *P < 0.05, **P < 0.01*. G: Western blot analysis of BCL-2, Bax, caspase-3 and β-actin expression in NP cells treated with HOT and siRNA-ITG α6. H: The results of western blot analysis are expressed as percentages of positive mean values ± SD. *n = 3, *P < 0.05, **P < 0.01*.

### ITG α6 protects against high oxygen tension-induced ROS production in NP cells

To investigate the effect of ITG α6 on HOT-induced ROS production in NP cells, 2’,7’-dichlorodihydrofluorescein diacetate (DCF-DA) staining was used to test the level of ROS. The fluorescence intensity of DCF-DA was observed with fluorescence microscope. The fluorescence intensity of DCF-DA was increased in NP cells exposed to HOT for 24h. Silencing of ITG α6 could further enhance the promoting effect of HOT on the production of ROS in NP cells (Figure 6A-B). We also applied flow cytometry to measure the fluorescence intensity of DCF-DA. The results showed that the peak fluorescence intensity of DCF-DA on flow cytometry was increased in NP cells exposed to HOT for 24h. Induction of ROS by HOT was further increased in ITG α6 silenced NP cells, indicating that ITG α6 could protect against HOT-induced ROS production in NP cells (Figure 6C).

**Figure 6.**
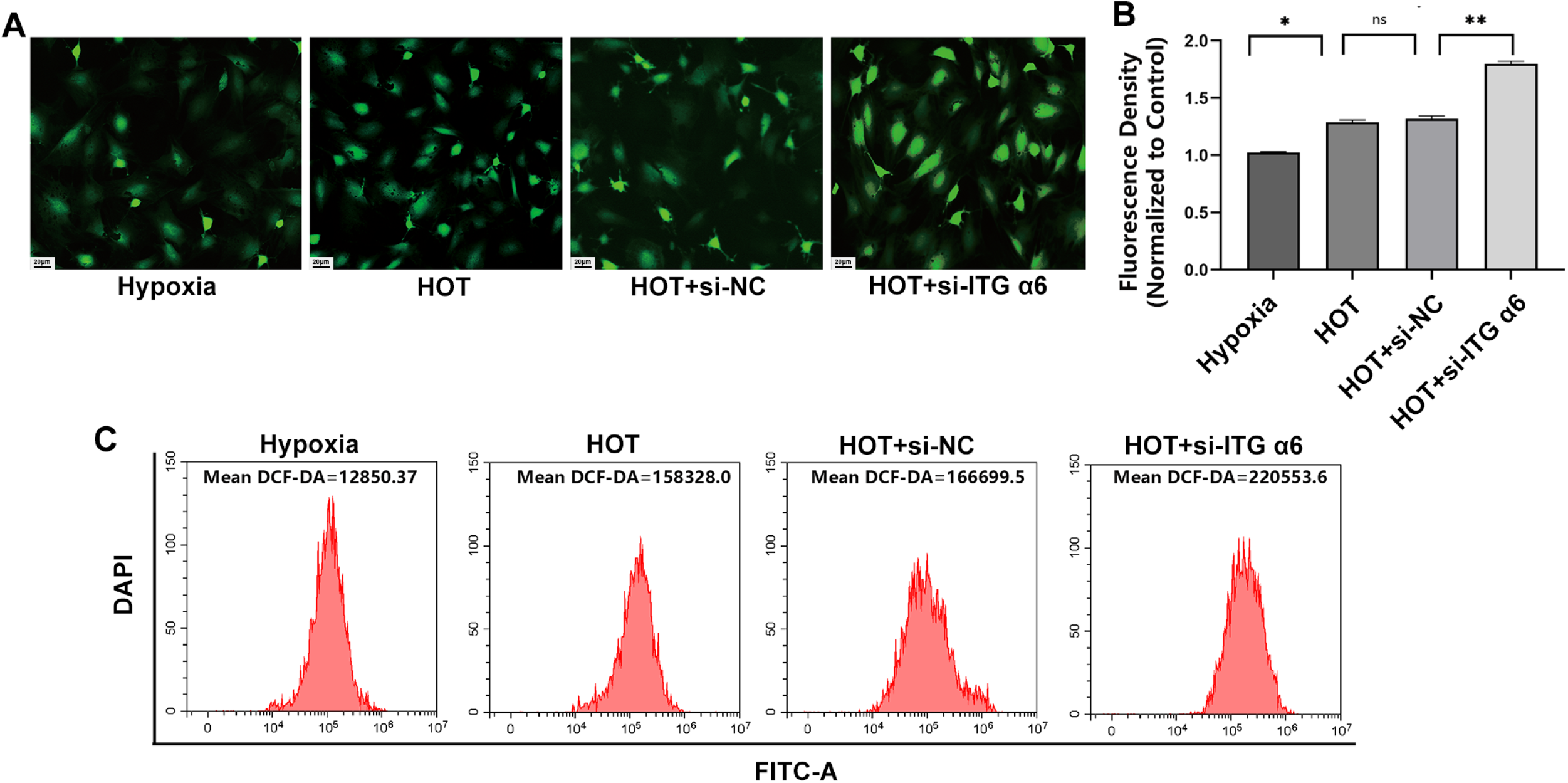
ITG α6 protects against high oxygen tension-induced ROS production in NP cells. A NP cells treated with HOT and siRNA-ITG α6 were stained with DCF-DA and detected by fluorescence microscope. B: Data were shown as mean ± SD. *n = 3, *P < 0.05, **P < 0.01*.C: NP cells treated with HOT and siRNA-ITG α6 were stained with DCF-DA and detected by flow cytometric analysis.

### The existence of an interaction and feedback regulation between HIF-1α and ITG α6 in NP cells exposed to HOT

The expression of HIF-1α was significantly decreased in NP cells exposed to HOT compared to cells cultured in hypoxia (Figure 7A-B). Silencing of ITG α6 obviously enhanced the inhibitory effect of HOT on HIF-1α expression, suggesting that ITG α6 could positively regulate the expression of HIF-1α (Figure 7 C-D). However, we found that silencing of HIF-1α could obviously upregulated ITG α6 expression, indicating that HIF-1α could negatively regulate the expression of ITG α6(Figure 7E-F). Similar results were also examined in HIF-1α knockout (HIF-1α ^-/-^) mouse model. The results of immunofluorescence staining showed that a strong red fluorescence signal of ITG α6 arose from the NP tissue of HIF-1α ^-/-^ mouse compared with WT mouse (Figure 7G-H). Based on the above results, we concluded that the increasing of ITG α6 expression by HOT contributed to maintain a NP tissue homeostasis through the interaction with HIF-1α.

**Figure 7.**
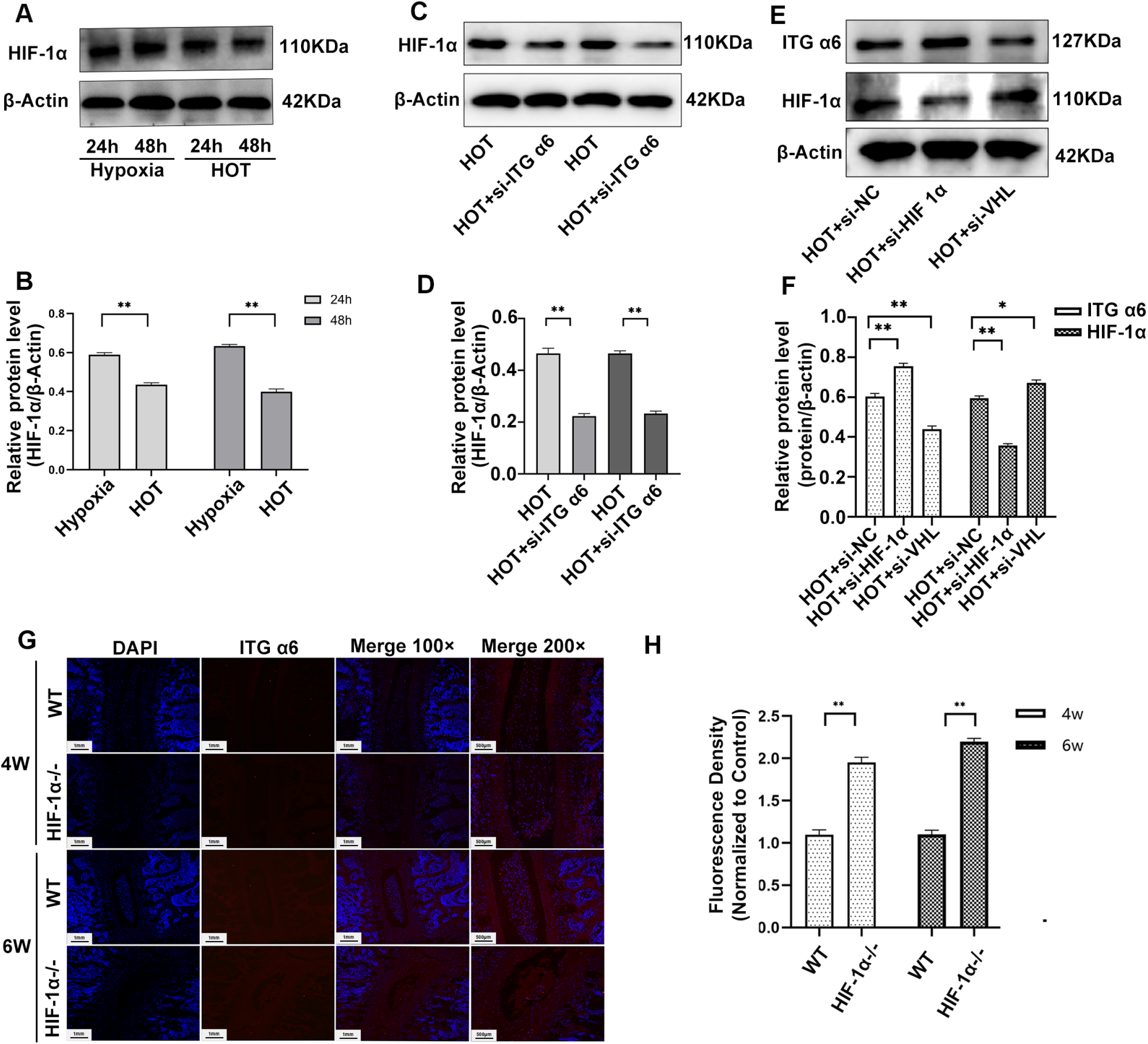
The existence of an interaction and feedback regulation between HIF-1α and ITG α6 in NP cells exposed to HOT. A: Western blot analysis of HIF-1α and β-actin expression in NP cells treated with HOT for 24h and 48h. B:The results of western blot analysis are expressed as percentages of positive mean values ± SD. *n = 3, *P < 0.05, **P < 0.01*.C: Western blot analysis of HIF-1α and β-actin expression in NP cells treated with siRNA-ITG α6 and HOT. D: The results of western blot analysis are expressed as percentages of positive mean values ± SD. *n = 3, *P < 0.05, **P < 0.01*.E: Western blot analysis of HIF-1α, ITG α6 and β-actin expression in NP cells treated with siRNA-HIF-1α or VHL and HOT. F: The results of western blot analysis are expressed as percentages of positive mean values ± SD*. n = 3, *P < 0.05, **P < 0.01*.G: Immunofluorescence staining of ITG α6 in knockout (HIF-1α ^-/-^) mouse model and WT mouse. H: Quantitative analysis of positive staining. The intensity of the immunofluorescence of ITG α6 was markedly increased in HIF-1α knockout (HIF-1α ^-/-^) mouse model. *n = 3, *P < 0.05, **P < 0.01*.

### Silencing of ITG α6 accelerates puncture-induced IVDD *in vivo*

To further determine the effects of ITG α6 on IVDD *in vivo*, siRNA ITG α6 nanoparticles (siRNA-ITG α6-CH), was injected into intervertebral disc to decrease ITG α6 expression *in vivo*. The inhibitory effects of siRNA-ITG α6-CH in NP tissue was examined by western blot. The results showed that the expression of ITG α6 was decreased in the NP tissues from rat injected with siRNA-ITG α6-CH compare with rat injected with siRNA-NC-CH (Figure 8A-B). To observe the effect of decreasing ITG α6 *in vivo* in rat puncture model, X-ray imaging was firstly used to determine changes in disc height index (DHI), the DHI was defined as the ratio of disc height to vertebral body height by lateral X-ray radiographs of caudal spine. X-ray imaging showed that the disc height of siRNA-ITG α6-CH rats was significantly lower than that of siRNA-NC-CH rats at 2 weeks after puncture (Figure C-D), indicative of IVDD. HE and Alcian Blue staining showed severe IVDD changes at 2 weeks after puncture in contrast to control group. The siRNA-ITG α6-CH group displayed significantly decreased NP content compared with the siRNA-NC-CH group at 2 weeks after puncture (Figure 8E-F). Col2α1 immunofluorescence also showed significantly decreased Col2α1 content in siRNA-ITG α6-CH group (Figure 8G-H). Taken together, these results demonstrated that ITG α6 might be a protection factor for IVDD.

**Figure 8.**
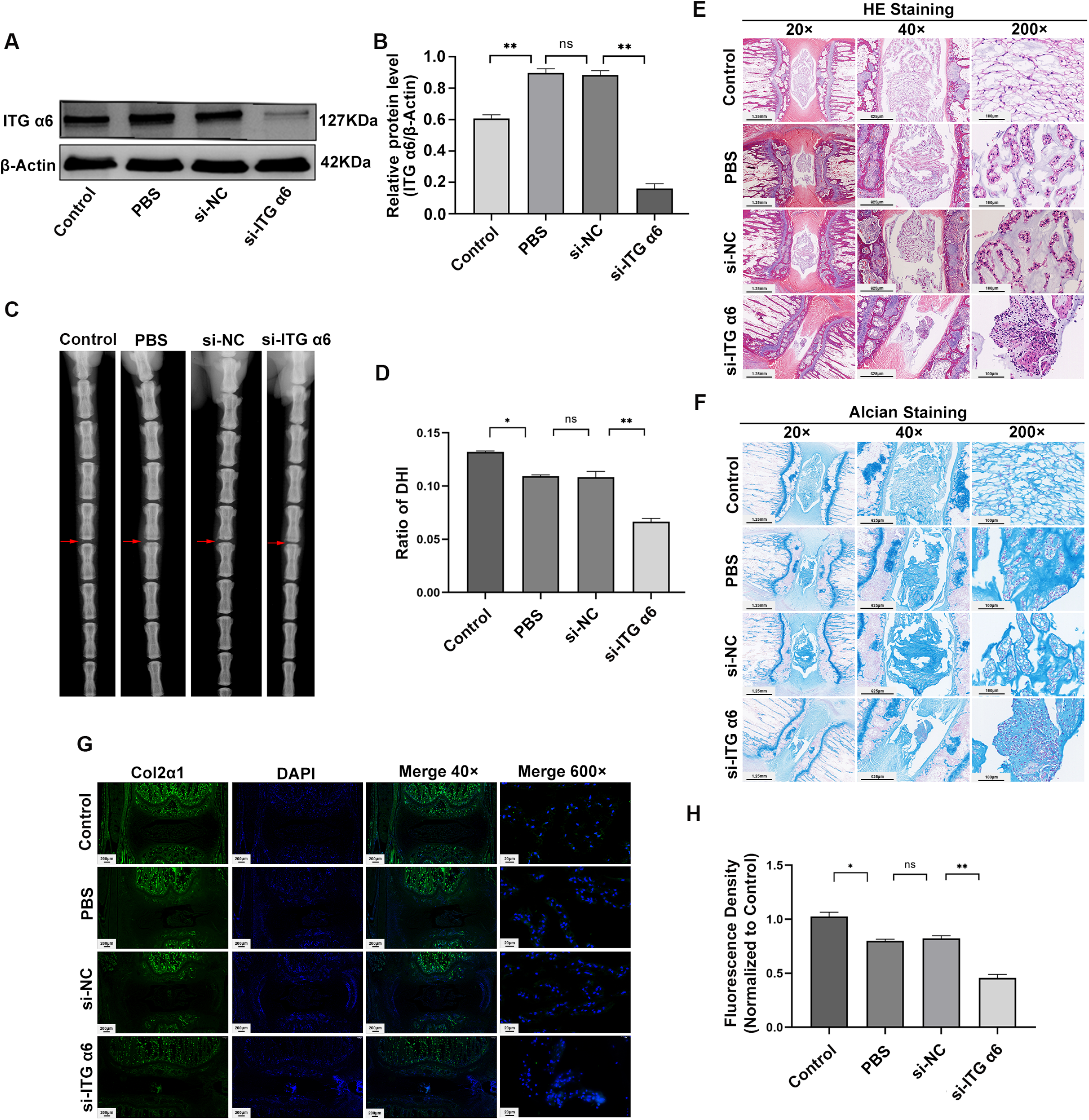
Silencing of ITG α6 accelerates puncture-induced IVDD *in vivo*. A: Western blot analysis of ITG α6 and β-actin expression of NP tissues in rat model treated with siRNA-ITG α6-CH. B: The results of western blot analysis are expressed as percentages of positive mean values ± SD. *n = 3, *P < 0.05, **P < 0.01*. siRNA-ITG α6-CH injected to IVD significantly decrease the expression of ITG α6 *in vivo*. C: X-ray imaging was used to investigate changes in disc height index (DHI) at 2 weeks after puncture compared with control group. D: The data of DHI were expressed as percentages of positive mean values ± SD. *n = 6, *P < 0.05, **P < 0.01*.E-F: HE and Alcian Blue staining showed severity of IVDD changes at 2 weeks after puncture compared with control group. G: Immunofluorescence staining of Col2α1 at 2 weeks after puncture compared with control group. H: Quantitative analysis of positive staining. The intensity of the immunofluorescence of Col2α1was markedly decreased in in rat model treated with siRNA-ITG α6-CH. *n = 3, *P < 0.05, **P < 0.01*.

## Discussion

Low back pain due to IVDD affect many individuals especially in the elderly population^2^. It often leads to the lower quality of life, which meanwhile results in direct and indirect annualized costs amounting to $50 billion in United States^29^. To identify the mechanisms of IVDD and find new therapeutic targets is imperative. The present study demonstrated for the first time that the expression of ITG α6 was significantly increased in IVDD at an early degeneration stage. The increasing of oxygen content of NP tissue was a critical factor for upregulation of ITG α6 in IVDD. Further studies found that ITG α6 could protect NP cells against HOT-induced apoptosis and ROS, and protect NP cells from HOT-inhibited ECM proteins synthesis. Upregulation of ITG α6 expression by HOT contributed to maintain a NP tissue homeostasis through the interaction with HIF-1α. Furthermore, silencing of ITG α6 *in vivo* could obviously accelerate puncture-induced IVDD. Taken together, these results revealed that the increasing of ITG α6 expression by HOT in NP cells might be a protective factor in IVD degeneration as well as restore NP cell function.

A latest study showed that 5% and 21% oxygen concentration led to decrease in anabolic gene expression of NPs while 1% led to increase, indicating that hypoxia was the most suitable and biological environment to maintain the NP phenotype and normoxia (21%, O_2_,defined as HOT in this study) was a pathological environment to NP cells^30^. The increasing of oxygen tension caused by abnormal neovascularization in discs is strongly related to the establishment and progression of IDD^31^. Currently, most studies focus on the protection effect of hypoxia on NPs^32–34^. However, very few studies were carried out to investigate the exquisite adaptation to oxygen stress of NP cells. In the present study, we explored which might be the most affected genes in ITG family by “adverse” oxygen conditions and might provide a potential target for treating IVDD. In addition, our results showed that HOT could increase the ROS production and inhibit ECM proteins synthesis. These results were consistent with previous studies, indicating that the cell model of HOT used in this study was successful and the results from these cell models were reasonable and reliable.

Many cytokines and chemokines have been confirmed to regulate cell metabolism by upregulating the expression levels of integrins^13^ ^35–36^. *Milica* et al. found that interleukin-8 stimulated trophoblast cell migration and invasion by increasing levels of MMP2, MMP9, ITG α5 and β1^35^. *Li* et al confirmed that CCL17 induced trophoblast migration and invasion by regulating matrix metalloproteinase and integrin expression in human first-trimester placenta^36^. However, no studies reported that the change of oxygen gradient could serve as a stimulator to regulate integrin expression and involve in the cell metabolism and growth. In the present study, we demonstrated for the first time that HOT increased the expression of ITG α6 in NP cells of IVDD. We further found that HOT upregulated ITG α6 expression by activating the PI3K/AKT signaling pathway, which played a key role in IVDD. In addition, we found that the level of ITG α6 was increased in the early stage of IVDD, whereas which was reduced in the middle and late stage of disc degeneration *in vivo*. It is likely that the upregulation of ITG α6 is a stress response of NP cells exposed to HOT in the early stage of degeneration. With the progress of IVDD, NP cells are subject to more severe microenvironmental stresses including starvation and oxygen stress due to the endplate calcification and metabolic waste accumulation^37^, resulting in severe destruction of NP tissue, which maybe one of the reasons for the low ITG α6 expression in the middle and late stage of disc degeneration. NP cells express HIF-1α, a key transcription factor that regulates anaerobic metabolism^38^. Unlike other skeletal cells, NP cells stably express HIF-1α in normoxia, while slightly increase induction in HIF-1α protein levels in hypoxia^39^. Peacock Brooks reported that expression of integrin α6 was positively regulated by HIF-1α in breast stem cells^40^. However, inconsistent with Peacock Brooks’s report from breast stem cells, ITG α6 was found to have an interaction and feedback regulation effect on HIF-1α in NP cells exposed to HOT. Silencing ITG α6 resulted in decrease in expression of HIF-1α. In reverse, silencing expression of HIF-1α led to elevated expression of ITG α6. Therefore, this might explain why HIF-1α was expressed constantly in normoxia in NP cells. Previous studies demonstrated that HIF-1α played an important role in NP cell’s metabolic regulation^39^. It promoted glycolytic pathway through increasing expression of numerous glycolytic enzymes, which is important for NP cells’ survival^41–42^. It is possible that increased expression of ITG α6 in NP cells in response to HOT contributes to maintain NP tissue homeostasis and NP cells’ survival through the interaction with HIF-1α.

The PI3K/ AKT signaling pathway, which regulates a wide range of cellular functions, is involved in the resistance response to hypoxia^43^. In this study, integrin α6 was also proved to be a downstream protein of PI3K/AKT signaling pathway, and it is in the opposite way as HIF-1α, influenced by the oxygen gradient. HOT activated the PI3K/AKT pathway, silencing PI3K/AKT signaling pathway would reduce the level of ITG α6. This indicated PI3K/AKT pathway was involved in the regulation of expression of ITG α6. HIF-1α is also regulated by PI3K/AKT pathway, however, expression of HIF-1α is elevated constantly in hypoxia, ITG α6 is elevated at early phase of HOT. Knocking out of HIF-1α resulted in increased integrin α6, this indicates both integrin α6 and HIF-1α regulated by PI3K/AKT pathway have a mutual effect on adjustment of NP cell’s metabolism, which protects cell from apoptosis, when hypoxia environment is lost.

There are several limitations of the present study. Firstly, although our results support that ITG α6 could ameliorate IVDD by regulating ROS production and apoptosis, however the particular downstream regulation mechanism remains unclear. Secondly, the underlying interaction between HIF-1α and ITG α6 need further elucidation. Lastly, the role of ITG α6 in IVDD require more detailed *in vivo* study.

In conclusion, the mechanisms of IVDD remain unknown and require further illumination. In the present study, we confirmed that the expression of ITG α6 was increased in the early stage of disc degeneration. The destruction of hypoxia environment in NP tissue is a leading cause of increased expression of ITG α6. Furthermore, we demonstrated that ITG α6 could protect NP cells against HOT-induced apoptosis and oxidative stress, and protect NP cells from HOT-inhibited ECM proteins synthesis. Silencing of ITG α6 *in vivo* obviously accelerates puncture-induced IVDD. Taken together, our study demonstrates for the first time that the ITG α6 is crucial protective factor for IVDD. These findings may provide new insights into the development of pharmacologic and physical therapies that can modify the course of IVDD.

## Acknowledgements

This work was supported by the National Natural Science Foundation of China, China [82072485] and the Shanghai Science and Technology Development Foundation [19ZR1456700].

## Ethics statement

Informed consent was given by the patients or relatives to obtain human intervertebral tissue at surgery. The experimental methods were carried out in accordance with approved guidelines and the study was authorized by the ethics committee of Changzheng Hospital, Naval Medical University.

## Author contributions

Zeng Xu performed experiments, analyzed data and wrote the manuscript; Jiancheng Zheng performed experiments and analyzed data; Ying Zhang and Huiqiao Wu analyzed data and approved the manuscript; Bin Sun and Ke Zhang performed the *in vivo* experiments; Jianxi Wang and Fazhi Zang collected human specimens and analyzed data; Xingkai Zhang designed the study, offered the HIF-1α knockout (HIF-1α ^-/-^) mouse model and analyzed data; Lei Guo and Xiaodong Wu designed study, analyzed data and wrote the manuscript.

## Data availability statement

The authors provide detailed description of methods and original data upon request.

